# Understanding the necessity of regulatory protein machinery in heterologous expression of class-III type of Ocins

**DOI:** 10.1101/2022.10.30.514460

**Authors:** Shilja Choyam, Rajagopal. Kammara

## Abstract

To date, there have been no or just a few reports of successful cloning and expression to create biologically active ocins or bacteriocins. Cloning, expression, and production of class I ocins are problematic because of their structural arrangements, coordinated functions, size, and posttranslational modifications. In the case of class III ocins, there are no reports of obtaining biological active proteins to date. Because of their growing importance, use and rapid functions require understanding mechanistic aspects to obtain biological active protein. As a result, we intend to clone and express the class III type. Also, by fusion or chimaera, they reshape class I types that lack posttranslational modifications into class III. Therefore, this construct resembles a class III type ocin. With the exception of Zoocin, expression of the proteins was found to be physiologically ineffective after cloning. But, few cell morphological changes such as elongation, aggregation, and the formation of terminal hyphae were observed. However, it was discovered that the target indicator had been altered to *Vibrio spp*. in a few. Finally, we confirm the existence of unidentified additional intrinsic factors for succesful expression to obtain biologically active protein.

## Introduction

One of the major attributes of probiotic microorganisms is the synthesis and production of ocins such as lactocin, bacteriocin, and enterocin that are antagonistic to closely related or unrelated organisms. The more specific activity of ocins makes them more rapid and specific in their action. There are reports stating the synthesis and secretion of ocins is a coordinated effort of the various regulatory regions that are located along the structural gene. Although ocins may be the best possible replacement for conventional antibiotics either for food preservation or for the generation of germ-free foods. They are preferred but not extensively used because of their low production, high cost, and lack of reproducibility. This situation prompts us to look for new ways to utilise heterologous expression and synthesis. The present manuscript deals with understanding the essentiality of the regulatory regions in heterologous expression. Therefore, we cloned and expressed a few ocins and succeeded in their expression in various *E. coli* hosts with and without regulatory regions. The manuscript looks into the possible factors playing a significant role in expression, leading to the obtaining of active protein.

Ocins are ribosomal synthesised antimicrobial proteins or peptides that kill or inhibit other related or unrelated bacteria. The presence of accessory proteins such as immunity proteins strengthens the host’s resistance to its own ocin. One source for ocins is from the original host and the other is from a heterologous host by recombinant expression. In most cases, the production from the native host is insufficient. Therefore, large-scale productions and industrial application are the need of the hour. In such situations, recombinant and heterologous expression are the major solutions. A few of the class II ocins have been successfully cloned and expressed in *E. coli* [1]. Synthetic production of ocins is another alternative, but the high cost of production makes it not feasible for commercial purposes.

Ocins are produced by both Gram-positive and Gram-negative bacteria. As per the recent ocin classification, the ocins of Gram-positive bacteria are divided into four classes. Class I bacteriocins (lantibiotics) are post-translationally modified to contain unusual amino acids. Nisin, subtilin, and variacin-8 are some of the examples. Non-lantibiotic bacteriocins are peptides that do not contain unusual amino acids and are heat stable. This class was further subdivided into three sub-classes: IIa (pediocin like bacteriocins), IIb (two-component bacteriocins) and IIc (thiol activated bacteriocins). Examples of class II bacteriocins include pediocin PA-1, lactacin F, etc. Bacteriocins of class III are high molecular weight, heat labile proteins that are either bacteriocidal or static [2].They are further subdivided into class IIIa (lysins) and class IIIb (non-lytic) examples, lysostaphin. The fourth class of bacteriocins, such as Plantaricin S and Leucocin S, are complex proteins with lipid and carbohydrate moieties. A combined method of the complete genome and peptidome of the bacteriocin-producer can be used to discover novel bacteriocins [3].

Ocins are capable of pore forming or inhibition of cell-wall, nucleic acid, or protein synthesis. They are cationic molecules, and they interact with the target bacterial cell surface, which is made of negatively charged lipopolysaccharides (LPS). While interacting, ocins are assigned in a configuration such that the positively charged groups interact with the negatively charged bacterial cell wall and the hydrophobic residues align together to traverse the lipid bilayer. This induces pore formation in the target cell membrane and eventually cell death occurs. A few ocins require receptors for docking with the target cell membrane, but some do not [4]. The effectiveness and efficacy of ocins action alone are heavily influenced by pH and temperature. A few ocins contain disulfide bonds and unusual amino acids that play a critical role in their activity [5].

Recent reports on the mechanistic aspects of the ocin bactofencin A [6] (using structure-function relationship and molecular dynamics simulations of ocin-membrane interactions) revealed that the C-terminus of the ocin is responsible for its activity and that the N-terminus is important in membrane interactions. The nature of amino acid residues in the C-terminal region determines the target specificity of the peptide. Many ocins have found promising applications in the areas of food preservation and antimicrobial therapy. The first and foremost requirement is the large-scale production of purified protein. The purification of ocin from the natural host, in most cases, was laborious, time-consuming, costly, not reproducible, and resulted in a low yield. The successful cloning and expression in *E. coli* increased ocin yield [7].

For example, unlike other proteins, the heterologous expression of ocins encounters many difficulties. The scale-up of ocins is low due to 1) the toxicity to hosts and 2) the multistep process of purification of ocins [8] (from fusion proteins, Wibowo and Zhao 2019). The existing reports conclude that nearly 30 % of currently approved recombinant therapeutic proteins use *E. coli* as their heterologous host. The well-established molecular biology, genetics, rapid growth and high yield production make *E. coli* a suitable host for hyper expression of recombinant proteins. Enterocins, the ocins produced by *E. faecium*, have antimicrobial activity against Gram-positive pathogens, especially *Listeria monocytogenes*. To facilitate biotechnological production of enterocins, an *E. coli* based expression system was used with fusion facilities for the purification [9].

The identification of ocins by public / free database mining has been promising, but their potential is difficult to evaluate in the absence of suitable expression systems [10, 11]. The success of heterologous expression of ocin primarily depends on the expression hosts. Subsequently, the nature of ocins, such as the presence of disulphide bonds, unusual amino acids, and post-translational modifications, affects its heterologous production. In some cases, the genes encoding immunity protein and transporter protein are necessarily incorporated for successful heterologous expression, leading to production. Finally, factors such as growth media, inducer agents, and temperature can all influence protein expression [12].

There are no standard rules in identifying and selecting expression hosts, plasmid characteristics, and fusion tags for the hyper-expression and production of large-scale functional ocins. Therefore, a trial and error method with the different combinations of vector and expression host with varying culture conditions is mandatory for the heterologous expression of ocins [13]. An expression system was developed for Class IIa bacteriocins which contain methionine, by making a fusion tag with thioredoxin. The high yield of ocin piscicolin 126 resulted from successful cleavage of the fusion tag [14]. In the present study, an attempt was made to over express selected high molecular weight (class-III type of ocins) ocins without any post-translational modification in the *E. coli* host. Even class I types were transformed into class III through chimaera formation and used. The rationale is that combining two or more bacteriocins can overcome this problem and broaden the activity spectrum [15].Therefore, various ocins from different sources were selected for the purpose, and different *E. coli* hosts were exploited for the heterologous expression.

Bacteriocins or antimicrobial peptides can be joined as dimers or pentameric bundles in various combinations to increase their activity and specificity, which is an advanced technology followed for large-scale production [16]. Many antimicrobial peptides have been generated for the eradication of *Streptococcus mutans*, the major causative agent of dental caries. The effectiveness of these antimicrobial peptides was further improved by synthesising them together as one molecule in various combinations (<u>17, 18, 19</u>). This combination was made by including a flexible peptide linker in between the antimicrobial peptides. The study concluded that the fusion peptides are lethal in comparison with original non-conjoined molecules [20]. A hybrid bacteriocin created by fusion of Laterosporulin and Thuricin S demonstrated greater antimicrobial activity than the individual bacteriocins, with a 55 % reduction in their MIC value against the indicator strain *S. aureus* [21]. The study was found to be interesting in showing the possibilities for the production of designer bacteriocins with a better killing ability [22]. A novel bacteriocin from Pseudomonas, PmnH, was identified with a dual toxin architecture [23]. It contains a colicin M domain, which interferes with peptidoglycan synthesis, and a colicin N-type domain with pore-forming ability. PmnH was considered as a natural chimeric bacteriocin with dual functions [24]. A recent study found that hybrid bacteriocins had higher antimicrobial activity and lower MICs than their individual counterparts [25]. A few studies by Acuna et al. showed that the hybrid bacteriocin Ent35-MccV showed antagonistic activity against both Gram-positive and Gram-negative bacteria [26]. Similar studies were reported, one among them was Chunhe et al. (2018) with a cecropin [27]. The second molecule being a chimeric bacteriocin of *Pyocin S* and *Colicin E*, observed to improve antimicrobial activity against *Pseudomonas aeruginosa* and *E. coli* [28]. The glycine linkers were incorporated at the gene level to increase the flexibility of the chimeric protein [29]. The special advantages of glycine, serine, and threonine linkers include rotational freedom of the polypeptide backbone, enhanced solubility, and resistance to proteolysis [30] with the intent of understanding the regulatory regions and their importance in occlusion expression. We exploit various ocins construct chimaeras and expressed in *E. coli* hosts with the objective of hyper expression and industrial scale production.

## Materials and methods Selection of ocins

A larger number of active bacteriocins from *Lactobacillus crustorum* were exploited by a combined genomic and proteomic approach [31]. Based on this study, ocins were selected according to their target organism, specifically food spoilage microorganisms. Selection criteria also include I) being a Class III type of ocin with a mol.wt range of approximately 10 kDa, and II) lacking post-translational modifications such as the presence of di-sulfide bonds and glycosylation sites. These attributes generally render ease of expression, and specifically food preservation and developing germ-free foods. Hence, we selected Linocin M18, Halocin H4, and Zoocin A. Other three class 1 ocins were chosen as chimaeras larger than 10 kDa. The brief details of the ocins selected for the study are given below:

### Linocin M18

Linocin M18 is an ocin produced by *Brevibacterium linens* M18 that has anti-listerial activity. The food industry has high demand for anti-listerial ocins. It was observed as a single copy gene in chromosomal DNA with 798 base pairs coding for 266 amino acids. Linocin M18 is an ocin that does not contain lanthionine [32].The Anti-Smash analysis of WGS concludes that it is a member of the bacteriocin family of proteins and contains core biosynthetic genes, additional biosynthetic genes, ABC-transporters, and regulatory and immunity protein genes.

### Halocin H4

Halocins are halo-archaeal equivalents of eubacterial ocins produced by *Haloferax mediteranei* and active against *Halobacterium salinarum*. The producer strain could grow in the presence of high salt conditions (4.0 M) and maintain intracellular salt concentrations of 0.3-2.0 M. Halocin adsorbs to sensitive cells, producing deformation and lysis and leaving empty ghosts in which the cell envelope seems to be intact [33]. The Halocin gene cluster was cloned and expressed [34]. It was found that the bacteriocin was produced in pro-form, which acts as a precursor for the mature peptide and finally undergoes post-translational modification to become a mature peptide. It is a plasmid-borne bacteriocin. As expected, the structure is similar to conventional ocins, where regulatory regions play a crucial role in their antimicrobial function.

### Zoocin A

Zoocin A is an ocin produced by *Streptococcus equi sub sp. zooepidemicus* antagonistic to *S. pyogenes* and *S. equisimilis*. Zoocin A was produced as a 285 amino acid pre-peptide that further undergoes cleavage of the leader sequence and produces a 262 amino acid active molecule. The estimated molecular weight of Zoocin is 31 kDa [35].

### Selection of class I type of ocins

In the present study, the peptide bacteriocins, which showed good antimicrobial activity, were selected for the construction of chimeric bacteriocins. Primers were constructed based on the sequence available in the NCBI GenBank and genomic DNA was isolated from *Lactobacillus* was used as the template to amplify the individual bacteriocins. Finally, by using splice overlap extension (SOE), a chimeric construct was made with the fusion of three individual bacteriocins.

### Construction of chimeric ocins

The construction of the various fusion/chimeric DNA was accomplished by the splice overlap extension (SOE) method [36]. The same method can be effectively utilised for site-directed mutagenesis [37, 38]. The method does not depend on restriction enzyme based scission and stitch [39]. In this method, two (or more) DNA fragments were joined together by employing PCR, without using either DNA restriction digestion or ligation [39]. The two fragments to be joined by SOE should have the highest possible complementary sequences at their respective junctions where they will join to form an “overlap”. These regions of complementarity are engineered into the DNA fragments, or blocks, to be joined (in the present case, two bacteriocin sequences) through separate PCRs, each employing primers specially designed. These two PCR-generated blocks are then used in the overlap extension in a third PCR, wherein the complementary strands hybridise partially at their 3’-ends through the regions of mutual complementarity after denaturation and reannealing. Thus, these two DNA strands act as mega-primers, and in the presence of thermostable DNA polymerase, the 3’-ends of this intermediate are extended to form a full length (i.e., fused), which may then be further amplified using the flanking primers derived from the first two PCRs used to generate the two DNA blocks. The hybrid bacteriocin poly-nucleotide fusions were made using three synthetic oligonucleotide PCR primers. Splice overlap extension PCR was carried out to (PCR III) obtain the hybrid DNA between ocin I and ocin II. In this reaction, approximately equivalent amounts of the purified DNA from PCR I and PCR II were mixed (representing approximately one fifteenth of the amplified DNA obtained from PCR I and PCR II) and carried out in a 100 µl reaction. To achieve optimal and targeted annealing between the hybridising areas of the two partially complementary strands from ocin I and ocin II. The reaction was first carried out in the absence of any other primer, using *Pfu polymerase* (Thermo Fisher Scientific) and the buffer, employing the following conditions: During denaturation (98 °C for 2 minutes), 4.0 minutes of RAMP were added to create slow cooling, which may aid in annealing complementary strands of both PCR products. For ten cycles, the same parameters were used to allow the formation of overlapped extended products. In the second phase, primers were added under hot start conditions for one more round of PCR. Another 25 cycles were given at the following conditions: denaturation (94 °C for 60 sec’s), annealing (40 °C for 60 sec’s), and extension (72 °C for 60 sec’s) to amplify the fusion products. Finally, after 1.0 minute at 72 °C to amplify the fusion product. At the end, after 10 min at 72 °C final extension, an aliquot from the PCR was analysed by agarose gel electrophoresis and purified through Thermo Scientific PCR purification columns. The relevant DNA cassettes were then purified by agarose gel electrophoresis, by excising the required DNA band and further purifying by the Thermo Scientific Gel Extraction kit. The full length extended, purified PCR products were kinased and then cloned into the pBluescript II KS-multipurpose cloning vector at the EcoR V site (which creates blunt ends). Screened the clones to identify the right recombinant containing ocin-ocin fusion product. Clones containing the SOE product were selected by restriction enzyme digestion to isolate the inserts and measure their size by agarose gel electrophoresis. The same procedure was followed for the fusion of the third ocin.

### Media, bacterial strains, and culture conditions

The strains used for the study are XL1Blue, BL21 (DE3), BL21 PlysS, and Origami (Novagen Inc., USA). The indicator strains used were Gram-positive and Gram-negative microorganisms. The strains were grown in Luria Bertani media (LB) (Hi-media Inc, Mumbai, India) with appropriate antibiotics as a selection marker. The indicator strains were grown on BHI media. The strains were cultured for 12 h at 37 °C and 200 rpm with commercially available pKS Bluescript II (Add Gene) and T7 expression systems (pET-23a plasmids, Novagen Inc, USA) were used for cloning studies.

### Preparation of chemical competent *E. coli*

To obtain a single isolated colony, streaked *E. coli* cells from glycerol stocks onto a fresh LB Amp plate and incubated at 37 °C for 12 hours. In 3.0 ml of fresh LB-medium, a single isolated colony was inoculated and incubated at 200 rpm at 37 °C. Subsequently, subcultured at 4.0 % of overnight culture to inoculate 100 ml of LB medium containing 10 mM MgCl_2_ and incubate at 37 °C until the absorbance at 600 nm is between 0.4-0.6. The cells were harvested by centrifugation at 4.0 °C at 4000 rpm for 10 min. Subsequently, they were re-suspended in 25 ml of ice-cold competent cell buffer (10 mM Hepes, 15 mM CaCl_2_, 55 mM MnCl_2_ and 250 mM Kcl), and incubated on ice for 10 min. Again, cells were harvested by centrifugation for 10 min at 4000 rpm at 4.0 °C. The procedure was repeated thrice to make the cells permeabilized. Finally, cells were dissolved in competent cell buffer containing Dimethyl Sulfoxide (DMSO). Subsequently, the cells were incubated at 14 °C for 10 min, and distributed as 100 µl aliquots in 1.5 ml tubes. The aliquots were subjected to a flash freeze in liquid nitrogen and stored at -80 °C for further use [40].

### Sub-cloning of ocins

Genomic DNA was isolated from the respective ocin producer strains by the GeneJET Genomic DNA Purification Kit (Thermo-Fisher Scientific USA). The primers for the specific ocins were constructed by incorporating restriction enzyme sites at 5’ and 3’ ends for cloning. Table 1 shows the list of primers used in the study. PCR amplification of ocins was done by *Taq DNA polymerase* (Thermo-Fisher Scientific USA). The initial denaturation starts with 94

**Table 1:**
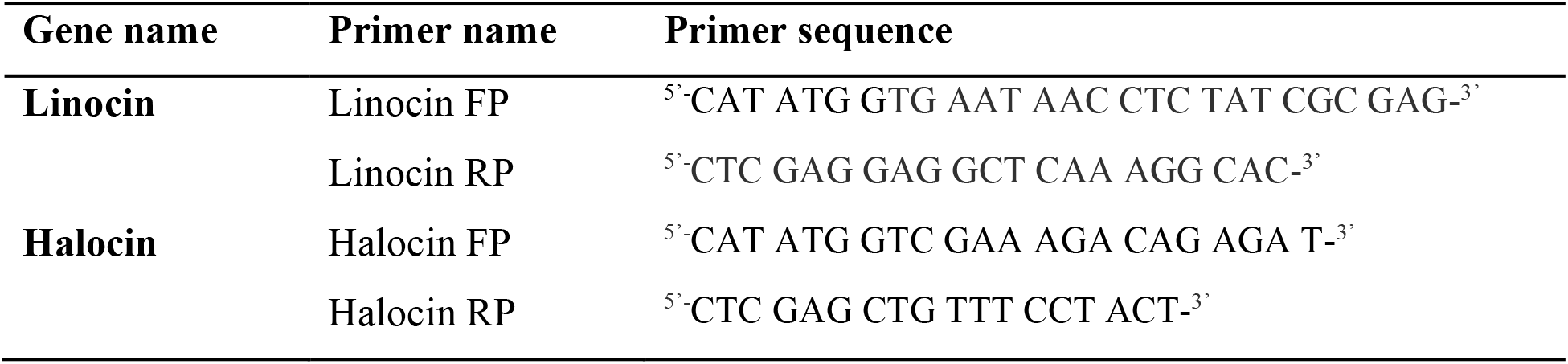
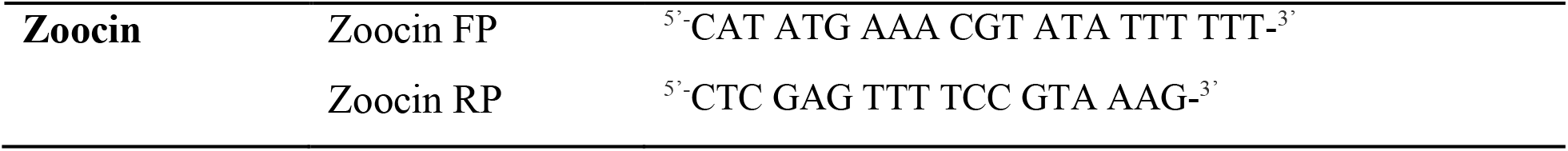
Primers used for the study.

°C for 5 min., a total of 35 cycles of 94 °C for 30 sec’s of denaturation, 48 °C to 52 °C for annealing, 72 °C for a 1.0 min extension, and a final extension at 72 °C for 5.0 min. The amplified product was subjected to purification by a PCR purification kit (Thermo-Fisher Scientific USA). The purified amplicon was cloned into the pBluescript II KS-cloning vector.

### Expression studies

Clones were transferred into expression hosts, BL21 (DE3), BL21 (DE3) plysS, Origami, and Rosetta (Novagen Inc., USA) for expression studies. The clones in different hosts were induced with various concentrations of IPTG to find an effective concentration. Different stages of growth were selected for induction, including 0.4-0.9 OD_600nm_ and 0.5-1.0 mM concentration of IPTG. After 3.0–4.0 hrs of induction, they were harvested, lysed with SDS sample buffer, boiled and subjected to tricine-SDS-PAGE analysis.

### Growth curve studies in various *E. coli* hosts

Clones transformed into various expression hosts, as well as control cells, were streaked on an LB-amp plate and incubated overnight at 37 °C. A single isolated colony from the plate was inoculated into fresh LB-amp. In media with an initial OD of 0.01 and continued to grow to a stationary phase, in between samples were taken for analysis. Clones and control cultures were incubated for 12 hours at 200 rpm at 37 °C. The growth was monitored by measuring OD at 600 nm at regular intervals. The growth curve was plotted as a graph of OD against time.

### Tricine SDS-PAGE

After induction, the resulting cells were harvested by centrifugation at 4000 rpm for 10 min. The harvested induced and un-induced cells were re-suspended in lysis buffer (150 mM NaCl and 100 mM tris with PMSF) and subjected to sonication. The sonicated cell lysate was further subjected to centrifugation at 12000 rpm and 4.0 °C for 30 min. The pellet and supernatant were processed for tricine SDS-PAGE along with a protein molecular weight marker, ran at 100 V for 2.0 hours [41]. The gel after electrophoresis was subjected to CBB (Coomassie Brilliant Blue) staining, and the protein bands were visualised under the white background.

### Western blot

Tricine SDS-PAGE of the protein samples, sonicated supernatant and pellet along with control were subjected to western blot analysis. The expressed proteins, after tricine SDS-PAGE, were transferred to a nitrocellulose membrane and probed with anti-His antibody. Finally, the developed blot with the substrate was analysed using the Gel-Doc Chem-doc gel imager (Azure Inc). The counterpart gel was CBB stained and compared with the western blot.

### SEM analysis

The effect of recombinant bacteriocin on the host cell wall integrity was analysed by SEM studies. Ocin clones from various indicator strains and a control (only vector) were streaked on LB-ampicillin plates and incubated for 12 hours at 37 °C. A single isolated colony from the plate was inoculated in LB-ampicillin broth and incubated for 12 hours at 37 °C, 200 rpm. Only 2.0 % of the overnight culture was used to subculture in 50 ml of fresh LB ampicillin broth following the same incubation conditions.

### SEM procedure

Induced and un-induced cells were harvested by centrifugation at 4000 rpm for 10 min, then the pellet was subjected to a series of alcohol washes, 10-100 % gradient. Finally, cells were re-suspended in the required amounts of absolute alcohol (50-100 µl). The pelleted cells were fixed in 2.0 % glutaraldehyde. Subsequently, they were incubated at 4.0 °C overnight. An aliquot of 2.0 µl of the sample was placed on a coverslip, desiccated, and observed under SEM. Clear and convincing images at 20x and 50x magnification were analysed.

### Well, diffusion assay

The culture supernatant and sonicated cell lysates were used to check the activity against the indicator strains. The well diffusion assay was followed to check the activity of the expressed ocin. The overnight grown culture of the indicator strain in BHI broth was mixed with BHI soft agar and poured over the agar plate. We allowed the culture to solidify by keeping it inside the laminar hood for 30 min, and subsequently wells were made in the soft agar plate. The sample was added into the well directly and allowed to diffuse into the medium completely before incubation. Once the sample inside the well had diffused completely, the plates were transferred into the incubator (37 °C for 12 h). The zone of inhibition in the plate was measured using the antibiotic zone measurement scale.

### Three-dimensional structure analysis of cloned bacteriocins

The predicted 3D model of the bacteriocin clone was analysed using the programme I-TASSER (Iterative Threading ASSEmbly Refinement) [42] for protein structure and function prediction. It automatically identifies and isolates reliable and identical structural templates from the PDB by the multiple threading approach LOMETS [43], with full-length atomic models constructed by iterative template-based fragment assembly simulations. The sequence was submitted to I-TASSER and analysed for three-dimensional structure using C-score [44].

The confidence of each model was quantitatively measured by a C-score that is calculated based on the significance of threading template alignments and the convergence parameters of the structure assembly simulations.

## Results

### Growth curve in different host strains

The OD (optical density) of the culture at 600 nm was taken regularly at an interval of 1.0 h for 12 hrs. Cultures were plotted against time. In Linocin, the growth rate was found to be decreasing when compared to the control in the expression host BL21 (DE3) and origami host strains. Growth rates were found to be normal in BL21 (DE3) pLysS (figure 1). Growth rates of Halocin were reduced in the host strains BL21 (DE3) and Origami, while an unaffected growth rate was observed in BL21 (DE3) pLysS host (figure 1). When Zoocin was expressed in BL21 (DE3) and origami hosts, a variation in growth pattern was observed wherein a delay in log phase and exponential phase were prominent. Their growth rates were normal when they were expressed in BL21 (DE3) pLysS (figure 1A to I). The growth rate of BL21 (DE3) hosts carrying chimeric molecules was less when compared to the control. A reduction in the log phase and exponential phase was observed. This may be due to the leaky expression of the protein in the latent stages of growth. When BL21 (DE3) pLysS host was considered, the growth rate was not decreased when compared to the control.

**Figure 1:**
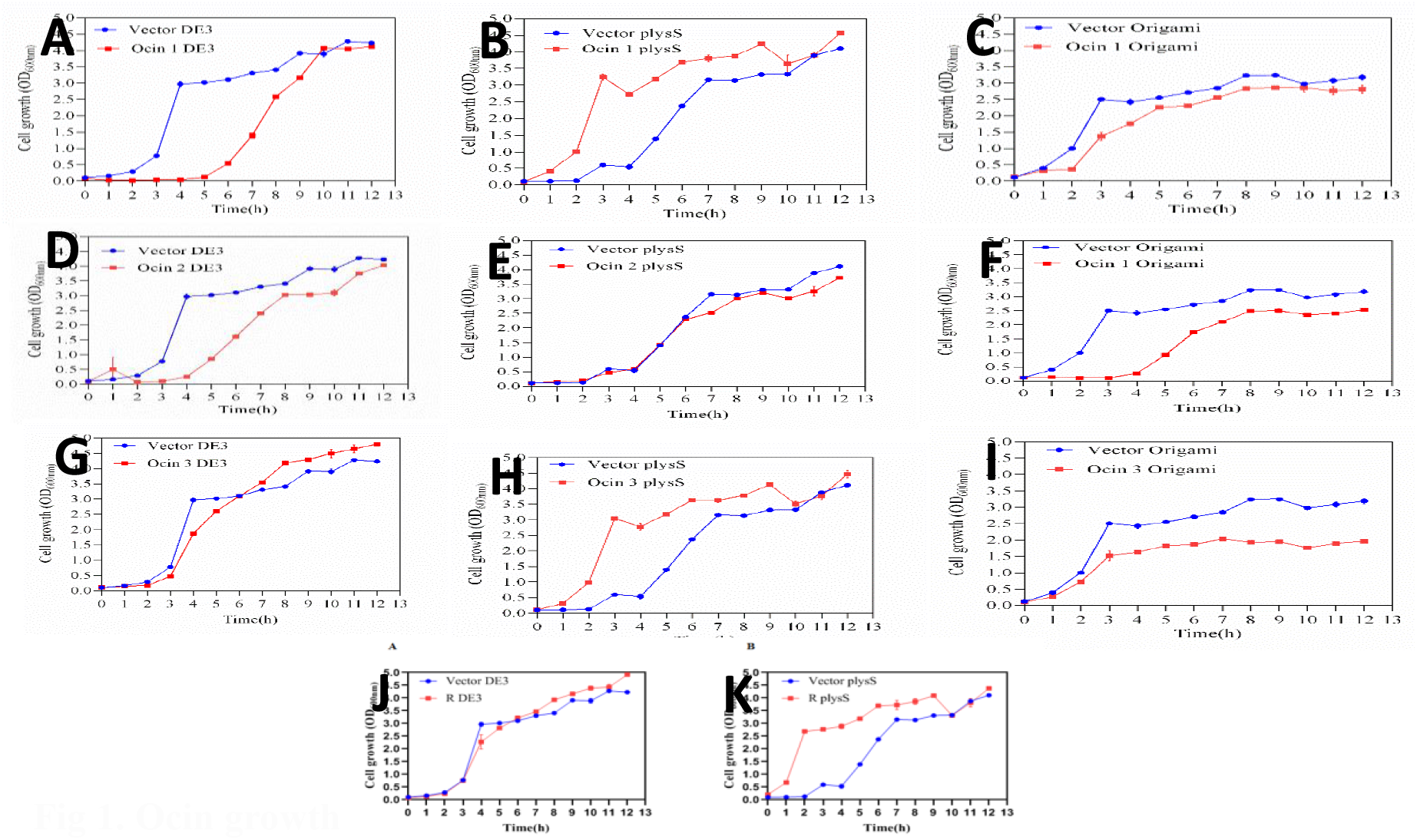
Ocin growth. Growth curves of Linocin, Halocin, Zoocin and chimeras/hybrids in BL21 (DE3), BL21 (DE3) Origami, and BL21 (DE3) pLysS. Procedure followed in brief: the single isolated colony was obtained by streaking and incubating on MRS Agar. A single isolated colony was inoculated into fresh MRS broth and grown at 37 °C 12-16 h shaking at 200 rpm. Grown till the OD reaches 0.5, later 2.0 % inoculum was sub-cultured in fresh MRS agar and OD was set at 0.01. Subsequently, grown at above conditions samples were collected every 30 min interval and OD _600 nm_ was checked at. Finally, graph was plotted between OD and time. A, B, C, D, E, F, G, H, I, J, and K. Each construct Linocin, Halocin, Zoocin, and hybrids were transformed into BL21 (DE3), BL21 (DE3) Origami, and BL21 (DE3) pLysS. Growth curves were done as explained. Vector denotes control BL21 (DE3).

### Chimeric ocins growth curve studies

The growth rate of BL21 (DE3) hosts carrying hybrid ocin was less when compared to the control. A reduction in the log phase and exponential phase was observed. This may be due to the leaky expression of the protein in the initial phases of growth. When BL21 (DE3) pLysS host was considered, the growth rate was not decreased. This may be due to the action of T7 lysozyme present in BL21 (DE3) pLysS host strains that hinders the overexpression of the cloned gene (also known as controlling the leaky expression phenomenon). The comparison of growth curves in BL21 (DE3) and BL21 (DE3) pLysS host strains in comparison with control is shown in figure 1.

### Protein expression in different host strains

It was discovered that ocin expression is highest when cells are un-induced and grown under normal conditions (37 °C, 200 rpm, for 12 hours). The suitable expression host for the maximum production of ocins was found to be BL21 (DE3). The induced culture showed a very low expression level compared with the un-induced one. The expression level of the chimeric ocin was high in BL21 (DE3) host strains when compared to BL21 (DE3) pLysS and Origami host strains. The various protein expressions are shown in figures 2A, B, C, and D.

**Figure 2:**
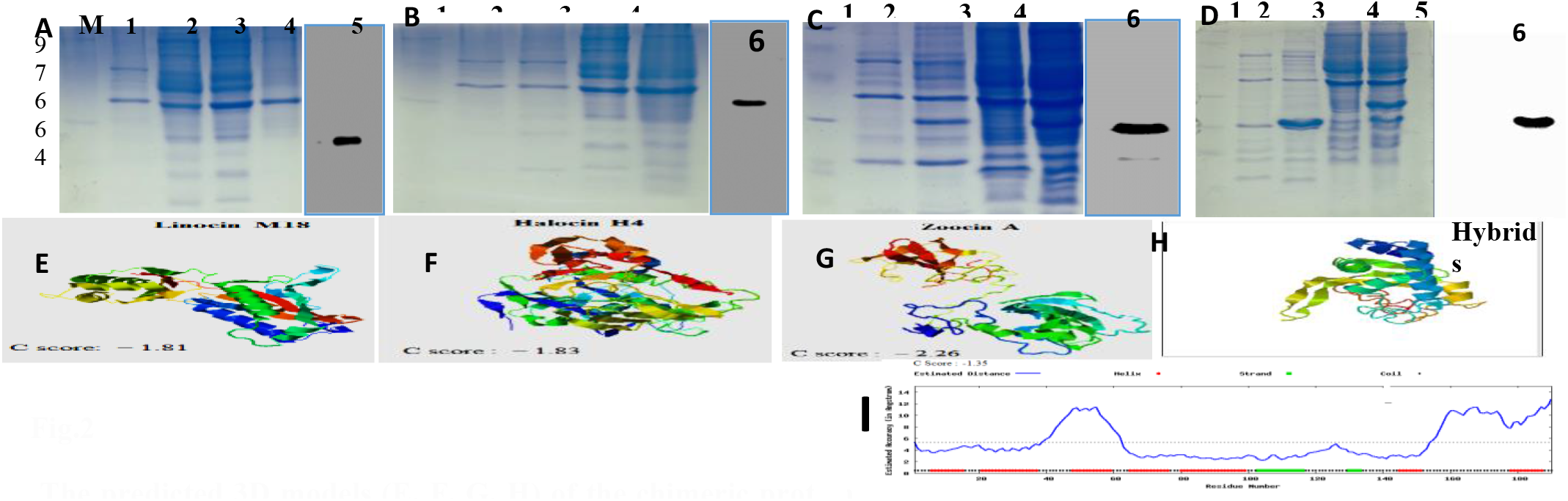
The predicted 3D models (E, F, G, H) of the chimeric protein with highest C-score and (B) estimated accuracy of the structure. Induction, expression, western blot and Secondary structure analysis of various ocins. The procedure in brief; after the bacterial growth overnight, 4.0 % inoculum was subcultured in fresh LB Amp, further grown to OD_600_ 0.4-0.5. Subsequently, (500 ml) cells were induced with 0.5 mM IPTG, growth was continued for 3.0 h. Cells were harvested by centrifugation at 9000 rpm for 15 min at cold conditions. The harvested cells were washed twice with cold ST buffer (NaCl 50 mM, and Tris.cl 150 mM pH 7.5) and re-suspended in 50 ml ST buffer. Sonication was followed at the following parameters 30 secs off, and 30 secs on for total 30 min. The sonicated cells were centrifuged to separate supernatant and the pellet. Only 100-200 µl of the lysate was processed for protein analysis, the remaining was subjected to Ni-NTA affinity chromatography. Out of which 20 µl cells were dissolved in 2x SDS sample buffer and boiled, centrifuged and loaded onto 12 % SDS-PAGE. The protein bands are shown in Fig. 2 A, B, C, and D. Fig 2A Lane 1: Marker, Lane 2: Linocin Un-Induced; Lane 3: Linocin Induced; Lane 4: Linocin Sonicated Pellet; Lane 5: Linocin Sonicated supernatant; Lane 6: Westernblot of Linocin; Fig. 2B. Lane 1: Marker; Lane 2: Halocin Un-induced whole cell lysate; Lane 3: Induced cell lysate; Lane 4: Sonicated pellet; Lane 5: Sonicated supernatant; Lane 6: Western blot probed with His-Tag antibodies. Fig. 2C. Lane 1: Molecular weight marker; Lane 2: Zoocin Un-induced whole cell lysate; Lane 3: Induced cell lysate; Lane 4: Sonicated pellet; Lane 5: Sonicated supernatant; Lane 6: Western blot probed with His-Tag antibodies **Western blot** Tricine SDS-PAGE of the protein samples, sonicated supernatant and pellet along with control was subjected for western blot analysis. The expressed protein after tricine SDS-PAGE was transferred to PVDF transfer membrane (mdi advanced micro-devices, Ambala, India) and further processed using Anti-6X His tag® and Goat Anti-mouse IgG H&L (HRP) antibodies (Abcam). Finally, developed the blot with western ECL substrate (BIO-RAD ClarityTM and Clarity MaxTM Western ECL Substrates) and analysed using BIO-RAD Chemidoc gel imager (ChemiDocTM XRS+system). The counterpart gel was subjected to CBB staining and compared with the blot. Identical procedures were followed for all ocins such as Zoocin, Halocin, Linocin and Hybrid ocins. **Three Dimensional structure analysis of Ocins** I-TASSER was used for the three-dimensional analysis of recombinant Ocins and chimeric protein. It is a hierarchical approach to protein structure and function prediction. It first identifies structural templates from the PDB by multiple threading approach LOMETS, with full-length atomic models constructed by iterative template-based fragment assembly simulations. The sequence was submitted to I-TASSER and analysed the three-dimensional structure using C-score. The confidence of each model is quantitatively measured by C-score which is calculated based on the significance of threading template alignments and the convergence parameters of the structure assembly simulations (Figure 2 E-I).

### Tricine SDS-PAGE and western blot

Tricine SDS-PAGE and western blot analysis showed the expression of cloned bacteriocins. All the ocins expressed were soluble in nature, as protein was observed in the supernatant and sparingly available in the pellet. Therefore, the protein is highly soluble. The results further confirm that the His-tag antibodies identify the presence of tagged ocins. The expressed chimeric protein was found in both supernatant and pellet but comparatively high in the soluble fraction. The various protein expressions are shown in figures 2A, B, C, and D.

### SEM image analysis

The SEM images of the host strains carrying bacteriocins revealed some morphological alterations when compared with the control. Elongation and distorted cells were observed in most of the host cells carrying ocin (Figure 3 A to D). Linocin protein expression in BL21 (DE3) host strains induced cell elongation, but not significantly in BL21 (DE3) pLysS hosts (Figure 3A). Aggregation and distorted cells were observed in the case of Origami™ B host cells after expression of the bacteriocin Linocin. Halocin expression also resulted in drastic elongation of cells in the BL21 (DE3) host and protein accumulation was observed in the case of BL21 (DE3) pLysS host strains (Figure 3B). Origami host cells were observed as aggregated with lysed cells after Halocin expression.

**Figure 3:**
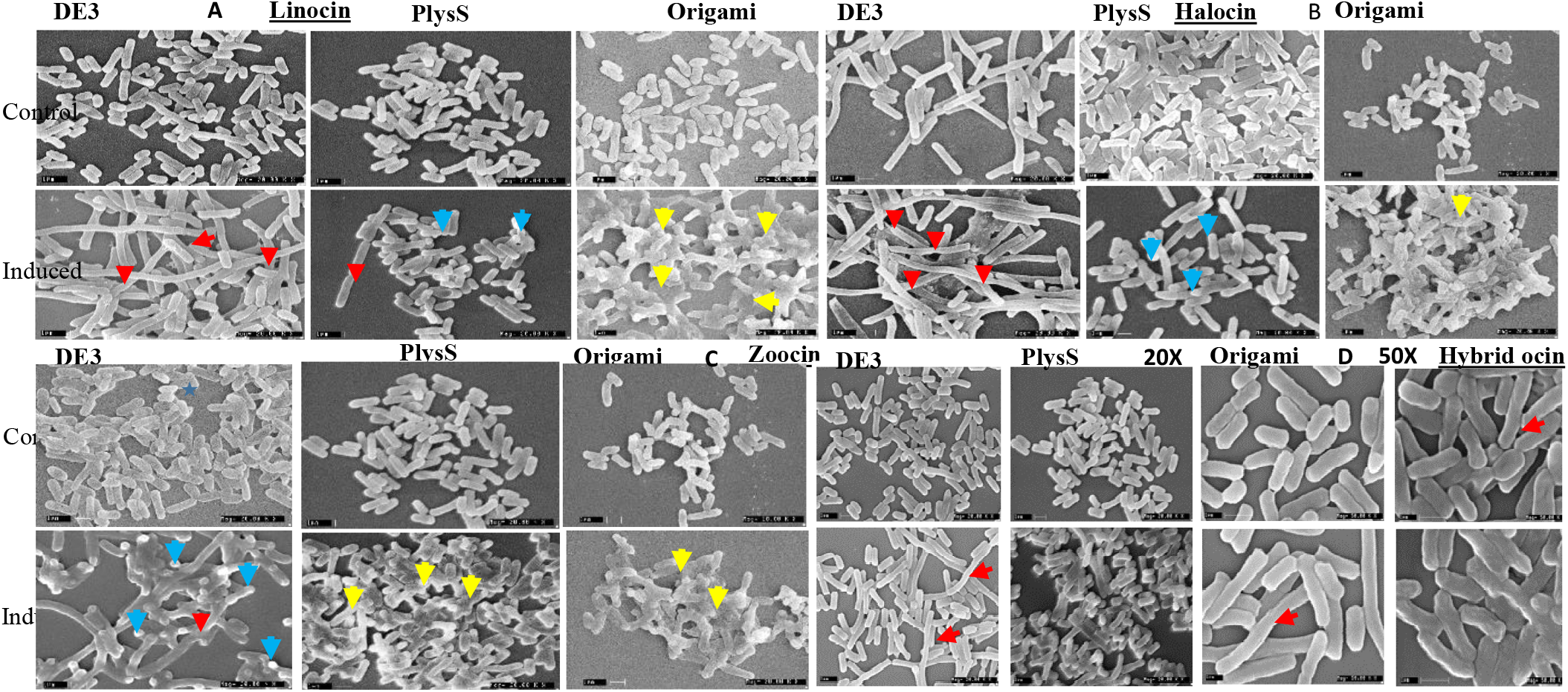
Scanning Electron Microscopy (**SEM)** studies The following procedure was followed to make samples for SEM studies: Bacterial cultures were prepared as mentioned in the growth curve studies. The subsequent overnight culture of 1.0 ml of sample was collected after each hour of incubation and processed for SEM. The cells were harvested and washed twice with phosphate buffer. The pelleted bacteria were fixed in 2.0 % glutaraldehyde and incubated at 4.0 °C overnight. After overnight incubation the cells were harvested by centrifugation at 4000 rpm for 10 min then the pellet was subjected to a series of alcohol washes, 10-100 % gradient. Finally, the cells were re-suspended in absolute alcohol (50-100 µL). An aliquot of ∼2.0 µL of the sample was placed on a coverslip, desiccated and observed under SEM. Clear and convincing images with 20x-50x magnification as per the requirements were analysed. Figure 3A represents Linocin expressed induced BL21 (DE3), pLysS, and Origami. Figure 3B. Represents Halocin expressed in induced BL21 (DE3), pLysS, and Origami. Figure 3C: Represents Zoocin expressed in induced BL21 (DE3), pLysS, and Origami; Figure 3D: Represents Hybrid ocin expressed in induced BL21 (DE3), pLysS, and Origami. 20x and 50x magnification images are shown. **Note:** Red Arrows indicate elongated *E. coli* host; Blue arrows indicate terminal hyphae; Yellow arrows indicates abnormal and aggregated cells.

Zoocin expression in all the host cells resulted in protein accumulation at the terminals. BL21 (DE3) host after Zoocin expression showed elongation and damage to the cell surface when compared with the control. Cells from BL21 (DE3) pLysS and Origami™ BL21 (DE3) cells also showed cell destruction and surface damage (Figure 3C). The expression of BAMP in the host resulted in protein accumulation inside the cell and elongated morphology when compared with control in the case of BL21 (DE3) and BL21 (DE3) pLysS host strains. The Origami host strain also underwent marked elongation upon expression of the cloned protein. The overall cell morphologies may be explained as BL21 (DE3) recombinants. They are observed to be elongated in a few instances and are abnormally elongated (shown in red). When the same constructs were expressed in BL21 (DE3) pLysS, no elongation structures were observed, but all showed dwarf structures, with the terminal hyphae being the recognisable result (Fig.3). Finally, the same construct when expressed in BL21 (DE3) Origami showed all cells abnormally aggregated. Therefore, this further states that there is an expression of cloned ocins that modifies their morphology.

It was observed that in the case of BL21 (DE3) host cells, the overexpression of the chimeric protein resulted in cell elongation (Fig.3A, Linocin, Halocin, Zoocin). The expressed chimeric protein may interact with the cell division process and prevent cell division, eventually leading to cell elongation. In the case of BL21 (DE3) pLysS host cells, the overexpressed protein resulted in cell destruction and the accumulation of proteins in the corner of the cell (Fig.3 A, B, C, and D). A white-coloured accumulation was observed in the terminal regions of the cells. The control cells were found intact without any morphological changes. The result of SEM analysis is shown in figure 3. Figure 3A represents SEM analysis of Linocin in BL21 (DE3), pLysS, and Origami. Figure 3B indicates the SEM analysis of Halocin in various *E. coli* strains. Figure 3C represents Zoocin, and Figure 3D represents hybrid ocin SEM studies. SEM analysis clearly states there is a compromised cell wall such as elongated cells, aggregated cells, and abnormal hyphae like terminal ends.

### Activity

The biological activity of the expressed ocins was found to be negative against most of the indicator strains. The recombinant Zoocin was active against the indicator strain *V. harveyi*. N-terminal amino acid sequence comparison with wildtype found that a stretch of particular amino acids might be responsible for the target-specific activity of the ocins. In the case of Linocin, the N-terminal and C-terminal changes occur for the production of active ocin by post-translational modification by cleavage. The cloned sequence and active protein sequence showed changes in some amino acids. In the case of Halocin, the same N-terminal and C-terminal modifications were observed. Zoocin A did not show any C-terminal modification but did show a deletion of amino acids at the N-terminal end. It was found that Zoocin A was expressed in an active form but the indicator specificity changed. This indicates that the presence of some specific amino acids in the N-terminal region has a role in the target specificity of the ocins.

The cell lysate and culture supernatant of the four bacteriocins cloned were taken for the antimicrobial activity assay by agar well diffusion. Among the four bacteriocins taken for the study, only Zoocin was expressed in an active form. The overexpressed Zoocin in the culture supernatant was found active against the indicator strain *Vibrio harveyi. Streptococcus* was the identified indicator for Zoocin in its native form. After cloning, the expressed protein showed a shift in the indicator strain from a Gram-positive *Streptococcus* to a Gram-negative *Vibrio harveyi*. The anti-listerial bacteriocin Linocin lost its activity against *Listeria monocytogenes* after heterologous expression. Halocin also gave negative results in well assay against the tested indicator strains. The cloned and over-expressed products of the novel bacteriocins, chimeric molecules, also failed to show activity against the indicator strains used. The results obtained by the well diffusion assay with over expressed protein against the Gram-positive and Gram-negative indicator strains are shown in table 2. None of the recombinant chimeric ocins were active against most of the indicators. Chimeric molecules were subjected to 3D-prediction analysis, and it was confirmed that they have helices and coils. Figure 2E-I shows the predicted structure.

**Table 2:** Ocins antagonistic activity against various indicators

| Sl No | Indicator organisms | Linocin | Halocin | Zoocin | Hybrid |
| --- | --- | --- | --- | --- | --- |
| **1** | *Pseudomonas putida* (-ve) | NA | NA | NA | NA |
| **2** | *Corynebacterium callunae* (+ve) | NA | NA | NA | NA |
| **3** | *Pseudomonas aeruginosa* (-ve) | NA | NA | NA | NA |
| **4** | *Salmonella typhi* (-ve) | NA | NA | NA | NA |
| **5** | *Enterococcus gallinarum* (+ve) | NA | NA | NA | NA |
| **6** | *Streptococcus thermophilus* (+ve) | NA | NA | NA | NA |
| **7** | *Staphylococcus aureus* (+ve) | NA | NA | NA | NA |
| **8** | *Vibrio harveyi* (-ve) | NA | NA | ++ | NA |
| **9** | *Streptococcus mutans* (+ve) | NA | NA | NA | NA |
| **10** | *Vibrio cholerae* (-ve) | NA | NA | NA | NA |
| **11** | *Listeria monocytogenes* (+ve) | NA | NA | NA | NA |

## Discussion

Since ocins are toxic to the producer strain, it is necessary to find a host that can tolerate the recombinant protein. Different *E. coli* host strains were chosen for the purpose. The experiment was conducted with and without IPTG induction. It was observed that *E. coli* can tolerate the recombinant protein inside the cell, but the level of expression was observed to be very low. In this case, the IPTG induction does not initiate or enhance the ocin expression. Similar results were obtained with *ColE1*-derived plasmids and different recombinants. There was no amplification or product formation upon IPTG induction. That may be due to the toxic effect of IPTG or strong inhibition of translation and chromosomal replication after the induction [45]. The expression levels were increased in BL21 (DE3) without IPTG induction. This further confirms the leaky expression, which is not harmful to the host; possibly the amount of ocin production is transient and low. This incident does not lead to an accumulation of ocin, therefore no toxicity.

The growth curve studies in different *E. coli* host strains showed growth inhibition in the initial stages when compared to the control. The production of the toxic protein may inhibit growth in the initial stages of growth. The morphological changes were observed when chimeric molecules were expressed. Elongated cells were observed, showing the possible effect of the ocins on interfering with cell division. That may be due to the influence of the increased ratio of product mRNA on cellular maintenance by reducing the synthesis of proteins necessary for growth and reproduction [46]. The SEM analysis of the recombinants exhibited clear morphological changes such as elongation and accumulation of proteins in the cytoplasm. The hybrid construct did not show any lytic activity towards the host strain. No physical parameters of the cell have been modified. In the case of the BL21 (DE3) host, a marked elongation of the cells was observed. This may be due to the inhibition of cell division by interfering with septa formation. Even though *E. coli* was considered as the preferable host for the production of bacteriocins, Enzymes and other proteins are sometimes expressed and recombinant production results in insoluble and non-functional proteins. This may be due to the lack of post-translational modifications, misfolding, aggregation, codon bias, and intracellular accumulation as inclusion bodies [47]. The non-functional hybrid bacteriocin may be due to the formation of misfolded ocin. The heterologous host cytosol is maintained in a reduced condition.

The ocins’ N-terminal amino acid sequence is critical to their activity. The changes in the amino acid sequence may alter the hydrophobicity and thereby affect the activity. A random mutagenesis study of pediocin AcH concluded that fourteen amino acid residues are important for the activity. One specific mutation changed the species-specific activity of the ocin [48]. Among the four ocins, cloned Zoocin was found to have antimicrobial activity against *Vibrio harveyi*, but it was supposed to be antagonistic to *S. pyogenes* and *S. equisimilis*. The change in target specificity may be due to the lack of post-translational modification in the heterologous host.

Most of the ocins, especially those from LAB, were synthesised as biologically inactive precursors or pre-peptides. This inactive form contains N-terminal extensions that will be cleaved off during their export and produce a biologically active form of the bacteriocin. If amino acid modification happens in the ocins, they come under class I lantibiotics and if not, they come under class II bacteriocins with unmodified amino acid [49]. This may be due to the lack of post-translational modifications, misfolding, aggregation, codon bias, and intracellular accumulation as inclusion bodies [47]. In the case of hybrid bacteriocin constructed in the present study, similar results were observed. The non-functional hybrid bacteriocin may be due to misfolded production inside the heterologous host. The loss of activity of the bacteriocin may be due to the instability of the proteins inside the cell or due to translational errors [50]. Similar results were observed when hybrids were constructed by fusion at the C-terminal of bacteriocins BacR1, Diversin V41, Enterocin P, Pediocin PA1, or Piscicolin 126 with the protein Intein-CBD and all the hybrid bacteriocins obtained were found to be inactive [1]. The C-terminal region of a few bacteriocins was found to be necessary for hybrid bacteriocin activity as well as specificity [51, 52].The hinge region of the bacteriocin also plays an important role in retaining the antimicrobial activity of the fusion constructs. The amino acids that allow flexibility in antimicrobial peptides are crucial for their antimicrobial activity [53].

It has been concluded that different ocins require different strategies for their heterologous expression with high levels of activity in an active form. In the present study, it was found that Zoocin A does not require immunity protein and secretion protein for its heterologous expression and activity. In the case of other ocins, it was found that the expression was possible in an *E. coli* host without IPTG induction, but the activity against the indicator strain was negative. This indicates that these ocins require post-translational modifications for their function. In some cases, post-translational modification of ocins may be used as a mechanism to control the activation of the toxicity of ocins, thereby providing a level of control and host immunity [54]. For the large-scale recombinant production of ocins, the host strain should be able to sustain the toxic protein without having any effect on growth and survival. One approach was to identify cloned ocin and host interactions for their growth and survival. It was observed that the recombinant ocin inside the host strain does not have a lethal effect but induces modifications leading to cell morphological changes such as elongation and bulged termini (Fig.3 A,B, C, D, E).

Heterologous expression is a common process followed globally for large scale production. The expression of protein requires regulatory machinery such as a ribosome binding site, promotor, start codon (ATG), and terminator. These are intrinsically available and fit into the commercially available expression vectors. Even the hosts have been developed as per the requirements. Hence, *E. coli* expression systems are well developed, and any protein may be easily and effortlessly expressed and produced. But, expression of bacteriocins, which are the products/secondary metabolites of *Lactobacillus* and *Bifidobacteria, is* successfully expressed in *E. coli* hosts. However, for whatever reason, the expressed protein lacks biological activity. The regulatory mechanism involved in bacteriocin expression requires structural genes, immunity proteins, and transporters, etc. When bacteriocin is expressed in *E. coli, the* activity attained is zero, mainly due to the lack of regulatory regions and the presence of post-translational modifications in ocin. In the present study, we have successfully cloned and expressed various ocins, even the hybrids. But the expressed protein does not show biological activity. The reason may be just the absence of regulatory regions as explained above. Zoocin, which is biologically active, does not require the unravelling of any regulatory regions.

The four bacteriocins, namely Linocin M18, Halocin H4, Zoocin A and chimera’s, were cloned and expressed under the T7 expression system. As the bacteriocins are known to be cytotoxic, different *E. coli* hosts were considered for the comparative study. Among them, the high level of protein expression was observed in BL21 (DE3), in comparison with BL21 (DE3) pLysS and Origami™. The maximum expression and no lethal effects were observed under normal growth conditions and without IPTG induction. Zoocin A was expressed in a functionally active form without any modifications. But, the indicator specificity of Zoocin A was changed from Gram-positive *Streptococcus* to Gram-negative *Vibrio harveyi* due to a change in a single amino acid in the N-terminal region. Due to the lack of post-translational modifications, the other three bacteriocins were expressed in a functionally inactive form. All the expressed bacteriocins negatively affected or compromised the surface morphology of the host strains. The study confirms that ocins are successfully expressed in heterologous hosts. No problems were observed, but it is essential to understand the factors that affect their biological activity once they are expressed. Out of four ocins studied, only one was biologically active, but others failed. Presently, we are involved in understanding the phenomenon in heterologous hosts. The most intriguing part may be that all the expressed ocins don’t contain posttranslational modifications such as disulphide bonds or glycosylation sites, yet they are not biologically active. The expressed chimeric protein induced abnormal elongation of the host BL21 (DE3) and terminal protein accumulation in BL21 (DE3) pLysS. The negligible or no activity of the chimeric bacteriocin against the selected Gram-negative and Gram-positive indicator strains may be due to the formation of non-functional proteins. The predicted 3-D structure of the chimeric product was also analysed using I-TASSER and the observed -helical structure with the hinge region (figure 2 I).

## Acknowledgements

MoFPI and LSRB are highly appreciated for their support in the form of grants in developing Ocins, Hybrid ocins. CSIR-UGC is highly appreciated for their funding support to Shilja Choyam. Finally we thank and appreciate CSIR-CFTRI for the facilities provided.

